# Biochemical Characterization of Emerging SARS-CoV-2 Nsp15 Endoribonuclease Variants

**DOI:** 10.1101/2022.05.10.491349

**Authors:** Isha M. Wilson, Meredith N. Frazier, Jian-Liang Li, Thomas A. Randall, Robin E. Stanley

## Abstract

Global sequencing efforts from the ongoing COVID-19 pandemic, caused by the novel coronavirus SARS-CoV-2, continue to provide insight into the evolution of the viral genome. Coronaviruses encode 16 nonstructural proteins, within the first two-thirds of their genome, that facilitate viral replication and transcription as well as evasion of the host immune response. However, many of these viral proteins remain understudied. Nsp15 is a uridine-specific endoribonuclease conserved across all coronaviruses. The nuclease activity of Nsp15 helps the virus evade triggering an innate immune response. Understanding how Nsp15 has changed over the course of the pandemic, and how mutations affect its RNA processing function, will provide insight into the evolution of an oligomerization-dependent endoribonuclease and inform drug design. In combination with previous structural data, bioinformatics analyses of 1.9+ million SARS-CoV-2 sequences revealed mutations across Nsp15’s three structured domains (N-terminal, Middle, EndoU). Selected Nsp15 variants were characterized biochemically and compared to wild type Nsp15. We found that mutations to important catalytic residues decreased cleavage activity but increased the hexamer/monomer ratio of the recombinant protein. Many of the highly prevalent variants we analyzed led to decreased nuclease activity as well as an increase in the inactive, monomeric form. Overall, our work establishes how Nsp15 variants seen in patient samples affect nuclease activity and oligomerization, providing insight into the effect of these variants *in vivo*.

## Introduction

Coronaviruses are a family of large, positive stranded RNA viruses that have known zoonotic potential [1]. The Middle Eastern Respiratory Syndrome (MERS-CoV) and Severe Acute Respiratory Syndrome (SARS-CoV-1) coronaviruses have caused several epidemics in the early 2000s. Currently a novel coronavirus, SARS-CoV-2, is responsible for a multi-year pandemic (COVID-19). Coronaviruses are also responsible for 10-30% of common colds [2]. RNA viruses, which replicate with an RNA-dependent-RNA polymerase are known to mutate frequently [3]. Mutations that confer a benefit to the virus are useful, but many mutations are neutral or deleterious. Evolution of RNA viruses is generally accepted to be driven by both genetic drift and negative selection (removing deleterious mutations) [4]. Coronaviruses encode proof-reading machinery that reduces mutational frequency, but it is still very high comparatively. Having low replication fidelity (i.e. a high error rate) allows coronaviruses to sample the mutational space by supporting a wide range of sequences during an infection, which leads to fast adaptation in response to selective pressures [5]. Mutations in viral RNA can also arise through damage or the action of RNA editors [6]. The enormous sequencing effort throughout the COVID-19 pandemic has presented the opportunity to analyze patterns of mutations and investigate the functional effects of predominant mutants [7–15].

The first two-thirds of the ~30 kb coronavirus genome encodes two open reading frames, pp1a and pp1ab, that encode 16 non-structural proteins (nsps) via a ribosomal frame-shifting mechanism [1, 16]. The latter one-third encodes structural and accessory proteins, such as the spike protein. Many of the nsps function in the replication transcription complex (RTC) and are necessary for viral replication [17]. Studies have shown mutational frequency varies across the SARS-CoV-2 genome [18]; mutation frequency seems to be higher in regions encoding structural proteins, and lower in the region encoding nsps [18, 19]. While structural proteins, especially the spike protein, have been extensively studied, nsps have had less focus [20–22]. However, given their conservation across coronaviruses and lower mutational rates compared to structural proteins, nsps make attractive anti-viral targets.

One such target is Nsp15, a uridine-specific endoribonuclease that processes viral RNA to prevent the activation of the dsRNA sensor MDA5 [23–25]. *In vivo* studies in animal models infected with coronaviruses with inactivated Nsp15 show lower mortality and reduced pathogenicity [23, 25–27]. Molecular studies of Nsp15 have revealed important details regarding how it processes RNA. SARS-CoV-2 Nsp15 is a hexamer formed from a dimer of trimers; monomeric Nsp15 is inactive [28–31]. Each protomer consists of an N-terminal (NTD), middle (MD), and catalytic endoribonuclease (EndoU) domain. EndoU domains are found in all kingdoms of life and share a catalytic triad with RNase A [32]. Like RNase A, Nsp15 uses a two-step transesterification mechanism to cleave RNA 3’of uridines [28]. Increased understanding of the molecular details of Nsp15 structure and function provides a platform to better understand the effect of mutations in the Nsp15 coding sequence. Interestingly, in certain variants, Nsp15 mutations have been noted as lineage defining markers [33]. For example, in an analysis of different Delta variant clades, Nsp15 H235Y is a marker for Delta clade C and K260R is a marker for Delta clade E.

Our goal was to analyze the mutations in Nsp15 from the first year of the COVID19 pandemic and characterize how mutations affect its endoribonuclease activity, either directly or through changes in its oligomerization state. We analyzed Nsp15 mutations found in sequences deposited in the GISAID database as of June 2021 [34]. We selected mutations from each domain (NTD, MD and EndoU) based on frequency of mutation as well as our knowledge of active site residues. We then carried out *in vitro* biochemical assays of the purified mutants to evaluate their oligomeric state and test nuclease activity compared to WT, in order to understand how these mutants seen in SARS-CoV-2 variants would impact Nsp15 function.

## Results

### Identifying Nsp15 mutants from SARS-CoV-2 genome sequencing

Approximately 1.9 million full length sequences of Nsp15 were extracted from the EpiCov section (allnuc0614) of the GISAID database, which is a worldwide repository of viral isolates. The sequence of Nsp15 of each viral isolate was compared to the original Wuhan isolate. In line with the relatively low replication fidelity of coronaviruses, single nucleotide variants were present across all three domains of the endoribonuclease. Of the full-length Nsp15 sequences (1038 bases), 1025 positions were observed to have nucleotide substitutions (Supplemental File 1), which were corresponding to changes in 341 out of 346 amino acids from Nsp15. However, 87% of these variants were observed in less than 100 isolates (Supplemental File 1). Six amino acid variants were observed in over 10,000 isolates, where five of them (N74N, 43,964; D79D, 15,684; L214L, 40,846; L217L, 74,888; N278N, 10,718) are synonymous and one (D220Y, 10,323) is non-synonymous (Figure 1A). For subsequent biochemical analysis we focused only on non-synonymous variants that led to a change in the protein sequence (Figure 1B). As expected, surface residues were more frequently mutated than core residues [19].

**Figure 1:**
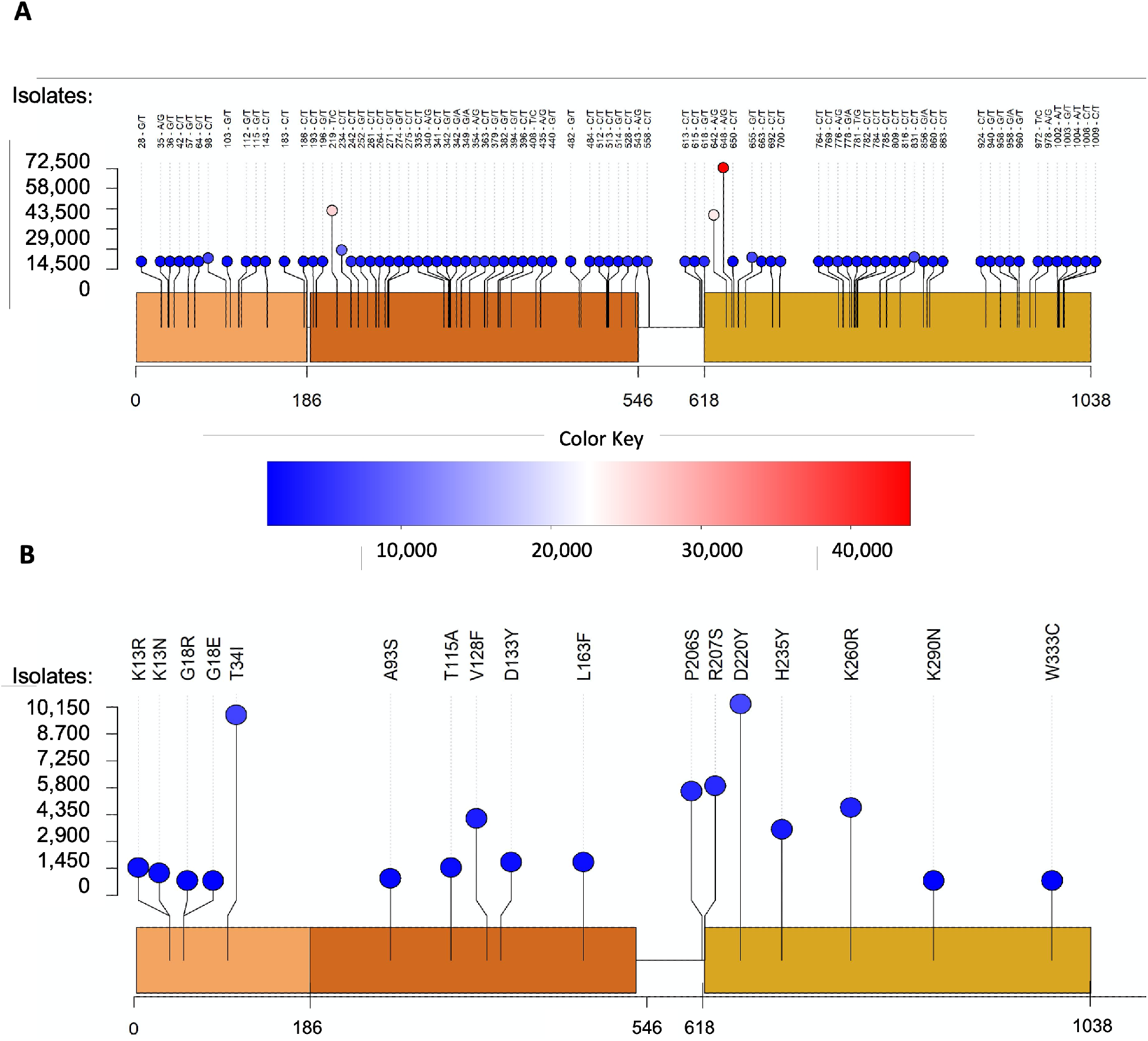
Variants identified in Nsp15 from the GISAID database. Approximately 1.9 million full-length NSP15 sequences (1038 bp) were downloaded from the GISAID database. The sequence of each Nsp15 isolate was compared to that of the original Wuhan isolate, and single nucleotide variants were then generated, which were presented in the lollipop-style plots. Each circle represents a variant observed at the indicated position. The height and the color of the circle indicate the frequency of the mutation event, ie. the number of isolates in which the variant was observed. **(A)** Variants are shown at the post-sequencing DNA level. Only variants found in at least 1000 isolates are shown. These single nucleotide variants were distributed widely throughout the three domains of Nsp15. **(B)** Selected variants are shown at the protein level. Variants with prominent numbers of isolates were selected to be biochemically characterized along with those found in residues previously shown to be important to the structure and function of Nsp15.

A total of 83 variants were observed in over 1,000 isolates which were unlikely to be sequencing artifacts. Among them, 53 variants are non-synonymous and 30 are synonymous. We selected several prominent mutations to study from these 53 non-synonymous variants: K13N (1716), K13R (1339), T34I (9900), A93S (1081), T115A (1810), V128F (4457), D133Y (1982), L163F (2072), P206S (5849), R207S (6174), D220Y (10323), H235Y (3630) and K260R (4938). In addition to these variants, others were chosen based on previous biochemical data [28, 29, 35, 36]. These variants were G18E (304), G18R (123), K290N (396), and W333C (304). Four of the selected residues (K13, T34, T115, and R207) had also been previously identified as major mutated residues of Nsp15 [18, 37]. The selected mutations covered all three domains of Nsp15 (Figure 2).

**Figure 2:**
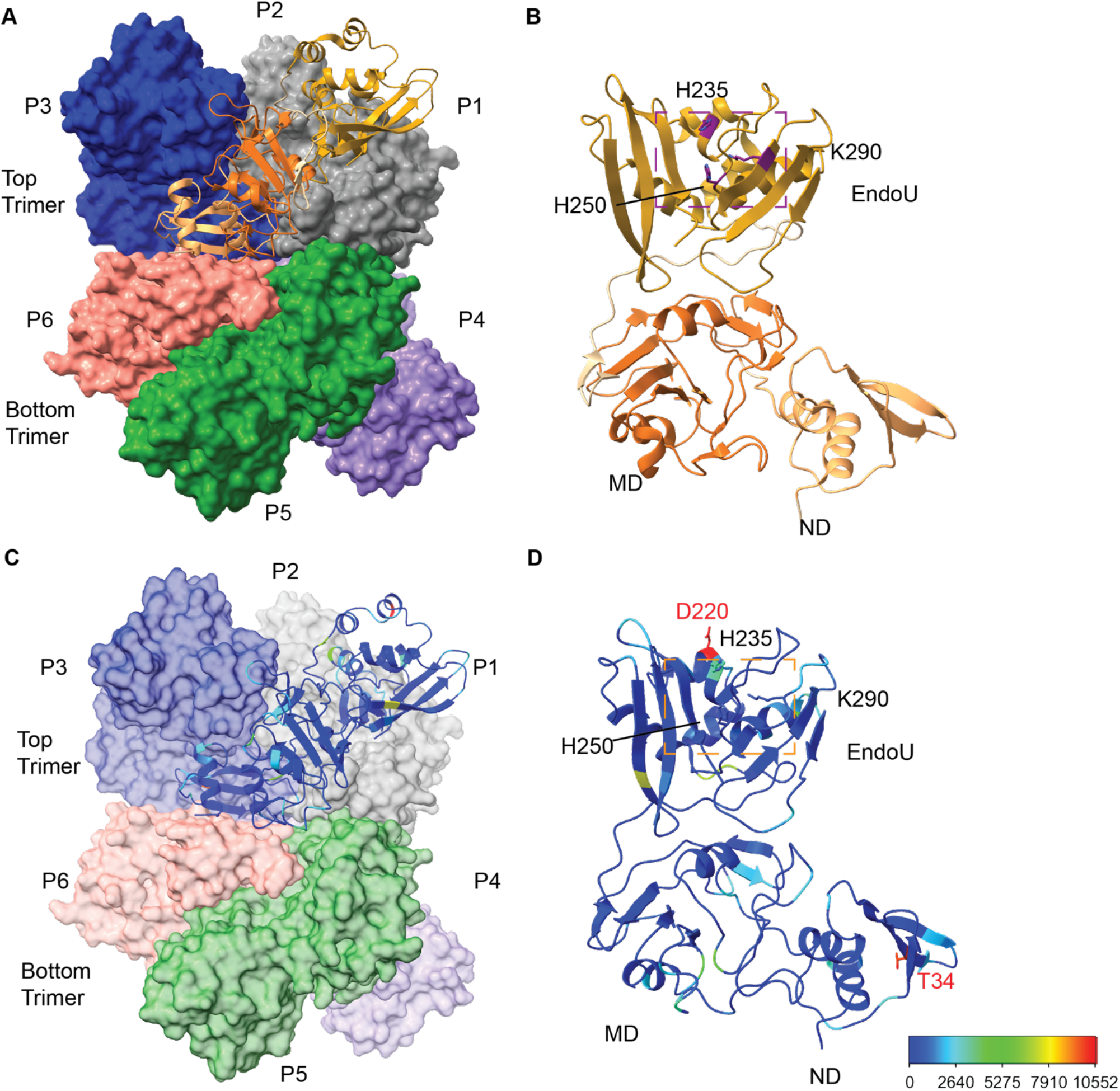
Overview of SARS-CoV-2 Nsp15 protein structure. **(A)** The Nsp15 hexamer forms from a dimer of back to back trimers. P1 is shown in ribbon diagram, while P2-6 are shown in surface representation (PDB ID: 7N06). **(B)** Zoom in of an Nsp15 protomer, colored as in Figure 1, by domain (ND, MD, EndoU). The catalytic triad is shown in stick representation, colored purple and labeled. **(C-D)** The number of mutations at each residue were colored using a rainbow palette (see scale bar at bottom left). (**C)** One colored protomer is docked into the hexamer. **(D)** Zoom in of the mutation mapped protomer. In addition to the catalytic triad, the two residues with the highest number of mutations are shown (T34 and D220).

Four of the Nsp15 non-synonymous substitutions show multiple base substitutions, G18R, R207S, K290N, and W333C. In each case there is a definite skewing of the abundance of one of these mutations. For G18R, the G to A substitution is 152-fold more prevalent than the G to C substitution. A simple explanation for this could be the elevated transition mutations relative to transversions, which was noted long ago [38]. However, the G to A transition on the positive strand of SARS CoV2 is one of the least frequent transition substitutions observed in SARS CoV2 [39]. The other three notable transversions are all either G to T or G to C transversions. The G to T transversion is 152-to 3087-fold more prevalent in these cases. This is substantially different from what is seen in either SARS CoV2 or human genomes [40] where G to C transversions are more common. The over-representation of the G to T events could also be due to a jackpot effect. While these observations were made with the GISAID allnuc0614 dataset (Supplemental File 1), these differences persist in newer versions (allnuc0215; Supplemental File 2). There has been significant recent analysis of genome-wide mutational spectra, especially as it relates to human cancer. Of particular interest has been the distribution of trinucleotide-centered mutational motifs and the mechanism(s) by which they occur. The only statistically significant trinucleotide motif found mutated in SARS CoV2 genomes was uCn [39]. None of the codons in the substitutions of interest within Nsp15 match that motif.

While the initial analysis of SARS-CoV-2 sequences in June 2021 provided the rationale for selecting Nsp15 mutations to characterize biochemically, we have continued to monitor SARS-CoV-2 variant lineages to understand how the Nsp15 mutations we selected appear in Delta and Omicron (Supplemental File 3). Delta, the predominant SARS CoV2 VOC through most of 2021, was replaced by the Omicron VOC beginning in late 2021. Omicron is not a direct descendant of delta; its origin is completely independent [41]. One can see this in our analysis of the mutational frequencies of the non-synonymous amino acid mutations within Nsp15 (Supplemental Table 2, Supplemental File 4). If Delta was the progenitor of Omicron, some of the mutations present in Delta would be present in nearly all of the GISAID Omicron genomes yet we see that for all non-synonymous substitutions only a fraction of the Omicron genomes carry any one substitution seen in the Delta genomes. Subsets of both Delta and Omicron genomes carry nearly all of these substitution mutations suggesting convergent evolution of these mutations in different SARS CoV2 lineages. Their persistence in both lineages suggests that any functional alteration to Nsp15 is well tolerated. It is also notable that the Delta genomes appear to have an almost uniformly higher mutation frequency for these substitutions. The higher mutation frequency noted for these substitutions in Delta lineage may also suggest a lower replication fidelity in this lineage, which could be of concern in terms of new Delta subvariants of concern emerging more rapidly from existing Delta strains.

### N-terminal domain variants

We assessed the impact of mutations on K13, G18, and T34I (Figure 3A) from the NTD of Nsp15, which has been shown to be critical for oligomerization [29, 37, 42]. In particular, K13 sits at the interface between neighboring protomers, and participates in water-mediated interactions with RNA ([29], Supplemental Figure 2). Given the importance of the NTD in oligomerization, we compared the amount of hexamer and monomer from each purified variant. WT Nsp15 has a hexamer/monomer ratio around 1 (Table 1, Figure 3B). K13R had a slightly greater amount of hexamer, while T34I had a ratio similar to WT. The other variants all had increased amounts of monomer. This provides additional evidence that disturbances in the NTD affect hexamer stability and suggests that these variants would result in less active Nsp15 in virus infected cells since monomeric Nsp15 is inactive.

**Figure 3:**
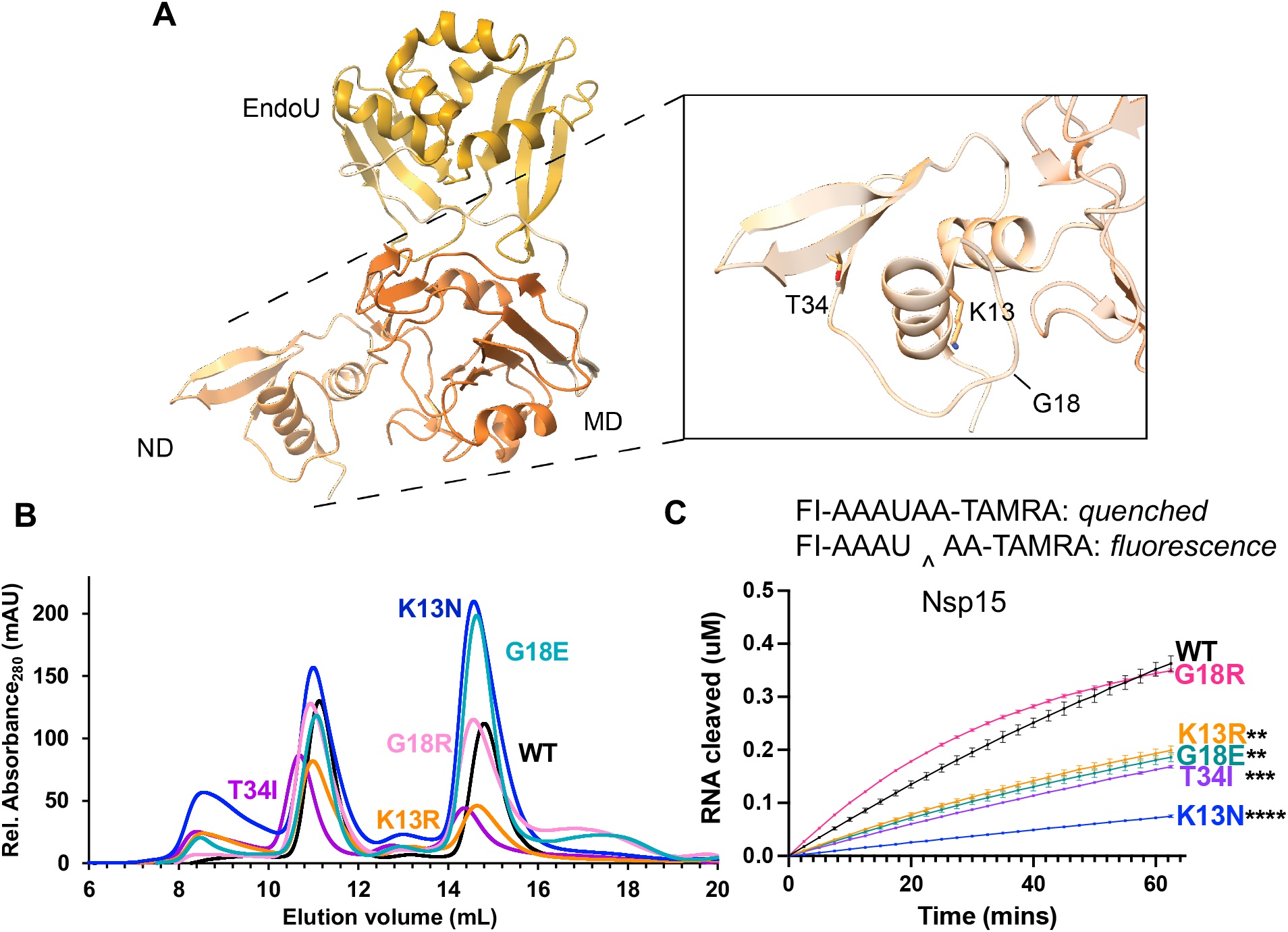
Characterization of Nsp15 N terminal domain variants. **(A)** Displayed on the structure are the residues K13, G18, and T34. **(B)** S200 elution profiles of NTD variants. Hexameric Nsp15 elutes at 11 mL, monomeric Nsp15 at 15 mL. **(C)** An embedded visual of the Fluorescence resonance energy transfer cleavage assay to accompany the FRET time course data for Nsp15 N terminal domain mutants. Nsp15 WT and variants (2.5 nM) were incubated with RNA (0.8 μM) at room temperature and fluorescence was monitored every 2.5 min for an hour. The average of a representative technical triplicate is plotted with standard deviation error bars. At least two biological replicates were performed for each mutant. Each mutant is represented by a different color: WT Nsp15 (black), Nsp15 G18R (pink), Nsp15 K13R (orange), Nsp15 G18E (teal), Nsp15 T34I (purple), and Nsp15 K13N (blue).

**Table 1:** Summary of Nsp15 variants analyzed in this study. A summary table of the variants categorized by domain of Nsp15. The rate refers to mutation rate as a percentage. The impact score *q* value describes its position in the distribution of all variants where the range is 0-1 and 0.5 is the mean q score[52]. The hexamer/monomer ratio is displayed along with the activity of the variant summarized by up and down arrows or NSC (no significant change).

| Residue | Rate (%) | Impact score (q) | Hex/Mono Ratio | Activity *** |
| --- | --- | --- | --- | --- |
| K13N | 0.070 | 0.83 | 0.63 | ↓ |
| K13R | 0.090 | 0.83 | 1.18 | ↓ |
| G18E | 0.006 | 0.24 | 0.56 | ↓ |
| G18R | 0.016 | 0.24 | 0.88 | NSC |
| T34I | 0.520 | 0.69 | 1.01 | ↓ |
| A93S | 0.057 | 0.30 | 0.49 | NSC |
| T115A | 0.095 | 0.14 | 0.55 | NSC |
| V128F | 0.234 | 0.36 | 0.50 | ↑ |
| D133Y | 0.104 | 0.73 | 0.48 | NSC |
| L163F | 0.109 | 0.33 | 0.23 | ↓ |
| P206S | 0.307 | 0.19 | 2.49 | NSC |
| R207S | 0.324 | 0.77 | 0.14 | ↓ |
| D220Y | 0.542 | 0.40 | 0.63 | NSC |
| H235Y | 0.191 | 0.90 | 1.61 | ↓ |
| K260R | 0.260 | 0.48 | 1.80 | ↓ |
| K290N | 0.021 | 0.61 | 2.12 | ↓ |
| W333C | 0.016 | 0.73 | 0.77 | ↓ |

Using an established FRET nuclease assay [28, 36], the cleavage activity of the hexameric forms of the K13N, K13R, G18E, G18R, and T34I variants was measured (Figure 3B). Nsp15 K13N showed the largest decrease in cleavage activity, while K13R, G18E, and T34I also had less activity compared to WT Nsp15. Nsp15 G18R nuclease activity was unchanged compared to WT Nsp15. Given the high number of isolates with T34I, its decrease in activity contradicts the notion that variants with high amounts of isolates are more advantageous for the endoribonuclease. T34 is a core residue in the NTD, surrounded by non-polar residues (Supplementary Figure 2). As a core residue, it is likely important for maintaining the correct fold of the NTD. While we would not expect the introduction of a non-polar residue to disturb the fold, the larger size of isoleucine may alter the NTD fold.

Overall, this FRET data supports the hypothesis that the NTD provides charge-mediated stability to RNA substrates [29]. The addition of a positive charge (G18R) did not negatively impact nuclease activity, while the loss of a positive residue (K13N) or the introduction of a negative residue (G18E) decreased activity. We also tested nuclease activity on a longer, more physiologically relevant RNA substrate corresponding to the Transcriptional Regulatory Sequence (TRS) for the SARS-CoV-2 N protein using a gel-based cleavage assay. The trends seen in the FRET assay were confirmed by gel-based cleavage (Table 1, Supplemental Figure 3).

### Middle domain variants

We analyzed middle domain variants from two distinct faces of Nsp15’s MD (Figure 4A). V128, D133, and L163 lie on one face of this domain, while A93 and T115 sit on the opposite face. All the MD variants had increased amounts of monomer, highlighting that the MD also plays an important role in stabilizing the hexamer. Previous work on MERS Nsp15 characterized several residues in the MD involved in oligomerization [35]. When we mapped these variants onto our SARS-CoV-2 Nsp15 structure, we see T115 and L163 facilitate interactions with neighboring protomers. T115 lies near the interface with a protomer from the bottom trimer, while L163 sits at the interface with a neighboring protomer in the same (top) trimer (Supplemental Figure 2). L163F dramatically reduced the amount of purified hexamer, suggesting this residue is particularly important for oligomerization (Table 1, Figure 4B).

**Figure 4:**
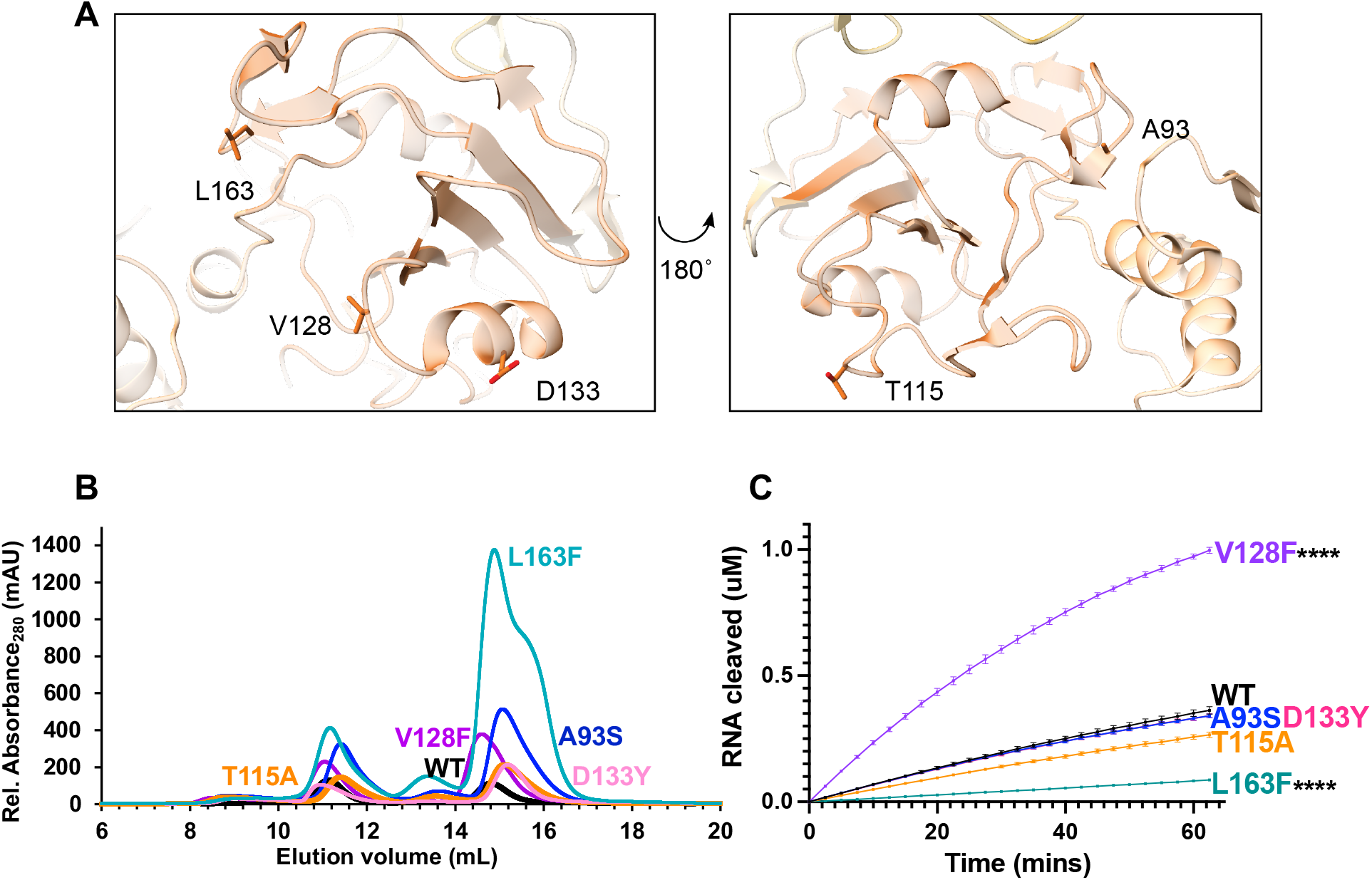
Characterization of Nsp15 Middle domain variants. **(A)** Displayed on the structure are the residues A93, T115, V128, D133, and L163. **(B)** S200 elution profiles of MD variants. Hexameric Nsp15 elutes at 11 mL, monomeric Nsp15 at 15 mL. **(C)** FRET time course data for Nsp15 Middle domain mutants. Nsp15 WT and variants (2.5 nM) were incubated with RNA (0.8 μM) at room temperature and fluorescence was monitored every 2.5 min for an hour. The average of a representative technical triplicate is plotted with standard deviation error bars. At least two biological replicates were performed for each mutant. Each mutant is represented by a different color: WT Nsp15 (black), Nsp15 D133Y (pink), Nsp15 T115A (orange), Nsp15 L163F (teal), Nsp15 V128F (purple), and Nsp15 A93S (blue).

Our FRET activity assay revealed a large, significant increase in activity with the V128F mutant, and a significant decrease in activity for L163F (Figure 4C). The remaining variants A93S, T115A, and D133Y showed no significant changes in cleavage activity. The activity trends seen by FRET were again confirmed using a longer RNA substrate in a gel-based cleavage assay (Table 1, Supplemental Figure 3-4).

Located in the middle domain, V128 resides in a surface exposed loop away from protomer interfaces; therefore, structural analysis does not provide a clear reason for this residue affecting cleavage significantly—Nsp15 V128F is two-fold more active than WT Nsp15. However, the change from a small hydrophobic residue to a large hydrophobic residue could possibly lead to changes in the domain conformation to avoid surface exposure. In virus, increased nuclease activity could potentially tip the balance from being protective (cleaving viral RNA to avoid dsRNA sensors) to harmful (preventing viral replication due to increased cleavage). Thus, our *in vitro* nuclease assays suggest this mutation could be detrimental to viral proliferation, yet it was found in numerous isolates. To better understand this mutant, studies in virus would be beneficial.

Along with a significant reduction in hexamer, the remaining L163F hexamer was less active than WT. The combined oligomerization and nuclease activity results for this variant suggest this mutant would produce very little active enzyme in virus. This residue is located near residues N164 in SARS-CoV-2 (equivalent to N157 in MERS, [35]). Mutation of MERS N157 decreased RNA binding affinity, reduced nuclease activity, decreased oligomerization, and decreased the thermal stability of the endoribonuclease [35]. Our analysis of L163F in SARS-CoV-2 supports earlier work that this interface is critical for nuclease activity and protein stability.

### EndoU domain variants

Finally, we analyzed specific variants from the EndoU domain, which contains the uridine binding pocket and catalytic core (Figure 5A). P206 (which exists at the end of the long linker between the MD and EndoU domains), R207, and K260 occupy the face of the EndoU domain opposite of the active site. H235, K290, and W333 lie in the active site on the other side of the EndoU domain and have been well-characterized biochemically [28, 29, 35, 36]. H235 and K290, along with H250, form the catalytic triad and interact with the scissile phosphate. It is therefore somewhat surprising that H235Y would be found in so many isolates (3630)—in fact, recent work characterized this mutation as a lineage marker for a clade of the Delta variant, Delta D. W333 forms important π-stacking interactions with the base 3’of the cleaved uridine. D220 occupies the same face as the active site but is farther removed from the RNA binding sites. It is surface exposed, and the side chain does not form any interactions with other residues (defined by a distance of < 5 Å). In contrast to the other domains, several EndoU variants showed an increased hexamer/monomer ratio: P206S, H235Y, K260R, and K290N. R207S, D220Y, and W333C showed a decreased hexamer/monomer ratio indicative of increased monomer (Figure 5B).

**Figure 5:**
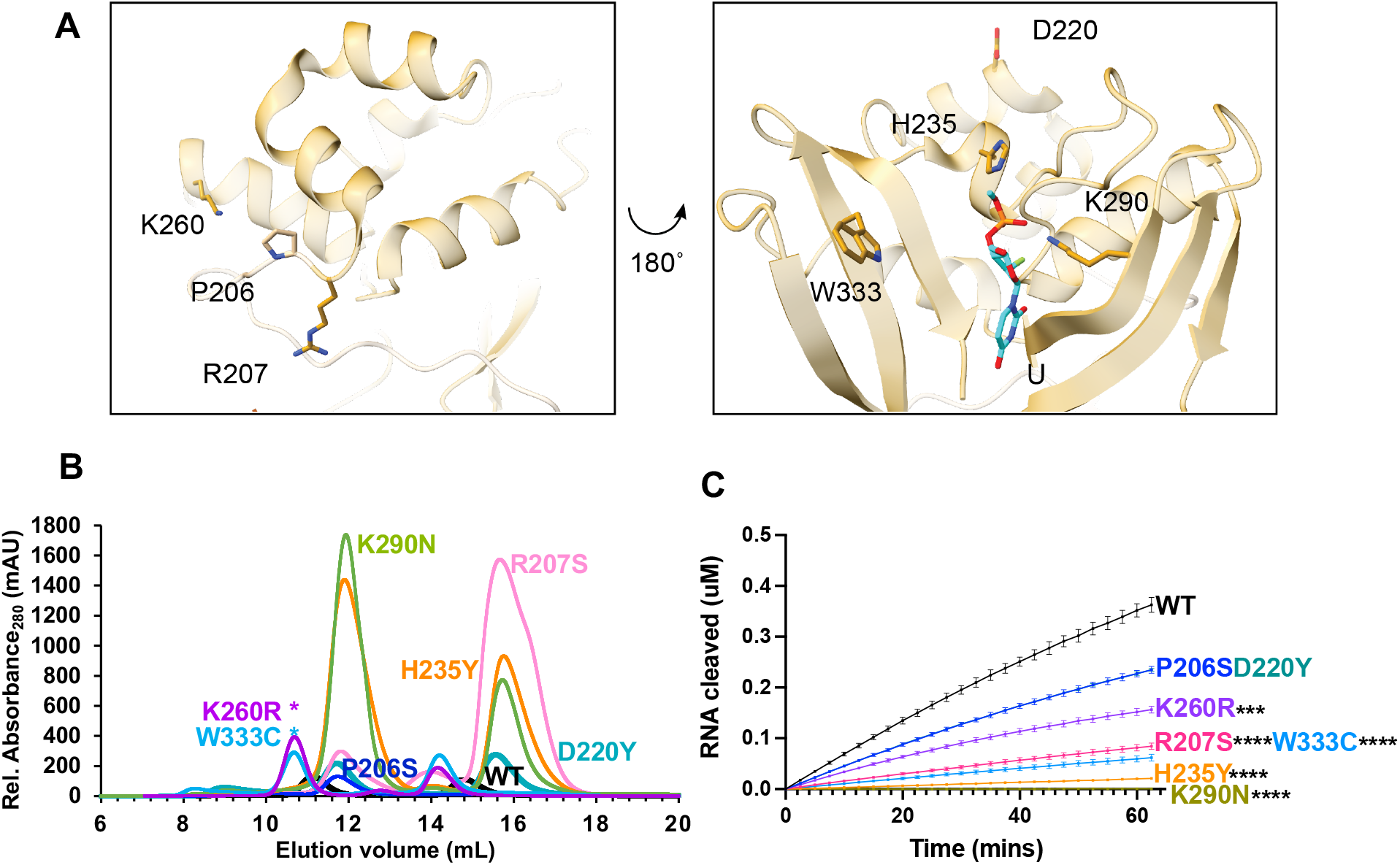
Characterization of Nsp15 EndoU domain variants. **(A)** Displayed on the structure are the residues P206, R207, D220, H235, K260, K290, and W333. H235 and K290 interact with the 3’-PO4 of uridine in the pre-cleavage RNA structure (PDB: 7N33) **(B)** S200 elution profiles of EndoU variants. Hexameric Nsp15 elutes at 11 mL, monomeric Nsp15 at 15 mL. **(C)** FRET time course data for Nsp15 EndoU domain mutants. Nsp15 WT and variants (2.5 nM) were incubated with RNA (0.8 μM) at room temperature and fluorescence was monitored every 2.5 min for an hour. The average of a representative technical triplicate is plotted with standard deviation error bars. At least two biological replicates were performed for each mutant. Each mutant is represented by a different color: WT Nsp15 (black), Nsp15 R207S (pink), Nsp15 H235Y (orange), Nsp15 D220Y (teal), Nsp15 K260R (purple), Nsp15 P206S (blue), Nsp15 K290N (green), and Nsp15 W333C (light blue). Asterisk (*) signifies mutants were separated using a different AKTA system which affected elution volume.

As expected, our FRET endoribonuclease assay confirmed that the catalytic residue variants H235Y and K290N were inactive (Figure 5C). The other active site residue W333C also showed significantly reduced cleavage activity, confirming its importance in pi-stacking with the RNA substrate. R207S and K260R also showed significantly reduced activity, while P206S and D220Y were not significantly different. Gel-based cleavage assays with a longer RNA substrate showed similar trends to the FRET assay data (Table 1, Supplemental Figure 5).

P206S and D220Y having similar activity as WT aligns with our initial hypothesis that mutants with high numbers of isolates would not significantly change nuclease function. Too much viral RNA cleavage and too little viral RNA cleavage are both likely to be detrimental to the virus, thus we would expect the more common Nsp15 variants to retain relatively the same nuclease activity.

## Discussion

In this work we biochemically characterized Nsp15 variants extracted from the GISAID database of SARS-CoV-2 sequences acquired during the pandemic. We specifically analyzed changes in oligomerization state and nuclease activity. The majority of the variants studied resulted in decreased Nsp15 activity. This work highlights the importance of combining bioinformatics and structural data to predict mutation effects and then testing those hypotheses *in vitro*. For example, based on Nsp15 structures alone, we would not have predicted a large impact of the T34I mutation, as it is a core residue that only interacts with other residues in the N-terminal domain. Indeed, the oligomeric state is not perturbed; however, our nuclease assays reveal a significant decrease in activity. Similarly, V128F would also not be predicted to have a great impact based solely on structural information, since it is not directly involved in any protomer interfaces and far from any EndoU active sites. However, our assays showed it decreases the formation of hexamer, but the hexamer formed has a substantial increase in nuclease activity. Biochemical characterization is not high throughput, therefore identification and selection of variants to test is important. Mutations with a high number of isolates are an easy selection criterion; our selection of K13, T34, T115, and R207 agree with bioinformatics analysis of main mutated residues across the SARS-CoV-2 proteome published in early 2021 [18]. Knowledge of molecular mechanisms is also important; this led us to select N-terminal residue and active site residue mutations to further test the established roles of these amino acids [29, 35, 36, 42]. We recently determined a structure of Nsp15 bound to dsRNA [43]; of the Nsp15 mutants we characterized here, the only non-active site residues in the area of the dsRNA are D133 (~5 Å) and V128 (~9 Å).

Coronavirus genomes have a bias for uridines [44, 45] so the evolution of a uridine-specific nuclease is interesting. Clearly, regulation of viral RNA through cleavage of uridines both in the polyU sequence of the negative strand (complementary to the polyA tail; [23]) and throughout the positive strand [46] is important for the coronavirus life cycle. Except for the Nsp15 H235Y mutation, which serves as a marker of a Delta viral clade [33], none of the active site residues accumulated mutations above the 1000 isolate threshold we used. The lack of nuclease dead isolates signifies the importance of nuclease activity for the virus. Thus, Nsp15 H235Y stands out as a candidate to further study in viruses.

A recent computational modeling study suggested Nsp15 serves as a scaffold for the assembly of the Replication Transcription Complex (RTC) [47]. In this function, nuclease activity may not be necessary. Thus, the H235Y and K290N mutations which kill catalytic activity but increase hexamer formation (Table 1), could support the RTC model. The Delta clade with Nsp15 H235Y has adaptations in other virally-encoded immune evasion proteins that may fill the gap of a nuclease dead Nsp15. However, a recent preprint analyzed transmission dynamics of SARS-CoV-2 variants and found the H235Y mutation along with several other mutations in that Delta clade led to transmission suppression compared to other lineages [48], evidence that a lack of Nsp15 activity may contribute to poor viral transmission.

Work to develop virus-like particles to study SARS-CoV-2 variants found that an important RNA packaging signal overlaps with the Nsp15 coding sequence [49]; therefore, variants that do not affect oligomerization or activity nonetheless could affect RNA packaging. Similarly, synonymous amino acid mutations may also result in changes to RNA packaging. More research is needed to understand the determinants of that *cis*-acting element [49]. Additionally, a recent preprint proposes an RTC model that positions Nsp15 to discriminate cleavage sites based on RNA structure as opposed to sequence [50]. This may be another way that variants impact function at the RNA level as opposed to the protein level.

Biochemical and *in vivo* studies of SARS-CoV-2 protein variants beyond the spike protein will contribute to a fuller picture of how the coronavirus proteome functions and changes over time. Recent work analyzing RdRp variants and the effect on remdesivir inhibition serve as one example [51]. Earlier this year a SARS-CoV-2 molecular dynamics database was created that takes the wealth of bioinformatics data and structural information we have on SARS-CoV-2 and applied molecular dynamics simulations to estimate the impact of variants [52].This high-throughput source of information is a great starting point to better understand how the virus has changed during the pandemic. The addition of *in vitro* data points, like the analyses presented in this paper, as well as *in vivo* analyses, to repositories like this database could facilitate a holistic understanding of the molecular impacts of viral mutations.

## Materials and Methods

### Nsp15 variants analysis

A total of 51,536,473 entries of the SARS-CoV-2 nucleotide sequences were downloaded from GISAID (version: allnuc0614). The downloaded data contains the coding sequences for genes including Nsp15 based on the hCoV-19/Wuhan/WIV04/2019 reference. Only full-length Nsp15 sequences (1038 bp) were retained for the downstream analysis, so 1,905,411 entries were included in the mutational analysis. The sequences of Nsp15 isolates were then aligned to that of the original Wuhan isolate (GenBank NC_045512.2) using the nucmer command from MUMmer 4.0 package [53] with default parameters. Variant data from the nucmer alignments were generated using show-snps command and the output was parsed directly using a custom perl script to convert MUMmer snps file to a tab delimited vcf-like table. Protein annotation was included in the final table (Supplemental File 1). Variants selected for testing were primarily at the amino acid level. Mutations that did not result in a specific amino acid change were discarded. The snps caused the same amino acid change (i.e. H235Y) were then summed to provide the total number of isolates per amino acid mutation. Visualizations of the variants on the Nsp15 DNA and protein sequences were generated via the lollipop function in trackViewer R package [54].

### Protein expression of Nsp15 variants

WT Nsp15 was previously synthesized by Genscript (Piscataway, NJ), and contains an N-terminal His tag with thrombin and TEV cleavage sites in pET14b [28]. For this study, Genscript mutated the WT sequence to our variants of interest (Supplemental Table 1). WT Nsp15 and Nsp15 variants were overexpressed in *E. Coli* C41 (DE3) competent cells cultured in Terrific Broth supplemented with 100 μg/mL ampicillin. Transformed cell cultures were grown to an optical density (600 nm) of 0.6-1.0 at 37°C prior to induction with 0.2 mM IPTG and overnight incubation at 16 °C. Harvested cells were stored at −80 °C until needed.

### Protein purification

Protein purification was carried out as previously described [28]. Cells were resuspended in lysis buffer (50 mM Tris pH 8.0, 500 mM NaCl, 5% glycerol, 5 mM β-ME, 5 mM imidazole) and then supplemented with complete EDTA-free protease inhibitor tablets (Roche). They were disrupted by sonication and the lysate was clarified by centrifugation at 15,000 rpm for 50 min at 4 °C, followed by incubation for 45 minutes with a TALON metal affinity resin (Clontech). His-Thrombin-TEV-Nsp15 variants were eluted with 250 mM imidazole and incubated with thrombin (Sigma) at room temperature in Thrombin Cleavage Buffer (50 mM Tris pH 8.0, 150 mM NaCl, 5% glycerol, 2 mM β-ME, 2 mM CaCl_2_) for a 4 h time period. Thrombin cleavage was quenched by the addition of 1 mM PMSF (phenylmethylsulfonyl fluoride). The cleavage reactions were incubated with TALON metal affinity resin, and tagless protein was eluted in batch and resolved by gel filtration using a Superdex-200 column equilibrated with SEC buffer (20 mM HEPES pH 7.5, 150 mM NaCl, 5 mM MnCl_2_, 5 mM β-ME). The peak fraction corresponding to the hexamer was used in subsequent assays. SDS-PAGE was used to assess protein purity (Supplemental Figure 1).

### Nsp15 endoribonuclease FRET assay

Real-time Nsp15 RNA cleavage was monitored as previously described [28, 29] with minor modifications to the protocol. The 5’-fluorescein (FI) label on the RNA substrate is quenched by its 3’-TAMRA label (5’-FI-AAAUAA-TAMRA-3’). A negative control substrate (5’-FI-AAAAAA-TAMRA-3’was also used (no cleavage observed; data not shown). The FRET RNA substrate (0.8 μM) was then incubated with a constant amount of Nsp15 variant (2.5 nM) in RNA cleavage buffer (20 mM HEPES pH 7.5, 75 mM NaCl, 5 mM MnCl_2_, 5 mM β-ME) at 25 °C for a 60 min time period. RNA cleavage was measured as an increase in fluorescein fluorescence. The fluorescence was measured every 2.5 min using a POLARstar Omega plate reader (BMG Labtech) set to excitation and emission wavelengths of 485±12 nm and 520 nm, respectively. Three technical replicates were performed to calculate the mean, standard deviation, and pairwise comparison test.

### Gel-based endoribonuclease assay

Gel-based cleavage assays were performed as described previously [29]. Double labeled RNA substrates (5’-FI and 3’-Cy5, 500 nM) were incubated with Nsp15 (50 nM) in RNA cleavage buffer (20 mM HEPES pH 7.5, 150 mM NaCl, 5 mM MnCl_2_, 5 mM DTT, 1 u/μl RNasin ribonuclease inhibitor) at room temperature for 30 min with the reaction samples collected at 0, 1, 5, 10 and 30 min. The reactions were quenched with 2x urea loading buffer (8 M urea, 20 mM Tris pH 8.0, 1 mM EDTA). Loading buffer without dye was used due to the expected size of cleavage products and the size of bromophenol blue. To monitor the gel front, control lanes of protein only with bromophenol blue were run. To generate a ladder, alkaline hydrolysis of the RNA was carried out for 15 min at 90°C using 1 μM RNA in alkaline hydrolysis buffer (50 mM sodium carbonate pH 9.2, 1 mM EDTA) and quenched with 2x urea loading buffer. The cleavage reactions were separated using 15% TBE-urea PAGE gels and visualized with a Typhoon RGB imager (Amersham) using Cy2 (λ_ex_ = 488 nm, λ_em_ = 515–535 nm) and Cy5 (λ_ex_ = 635 nm, λ_em_ = 655–685 nm) channels.

## Acknowledgements

We would like to thank Drs. Andrea Kaminski and Zachary Kockler for their critical review of this manuscript, and Dr. Juno Krahn for writing a Perl script converting number of mutations to PDB B-factors. This work was supported by the US National Institutes of Health (NIH) Intramural Research Program; US National Institute of Environmental Health Sciences (NIEHS, NIEHS/NIH ZIA ES103247 (RES) from the Division of Intramural Research of the NIH/NIEHS. This work was also supported by the NIH Intramural Targeted Anti-COVID-19 (ITAC) Program funded by the National Institute of Allergy and Infectious Diseases (NIAID, NIAID/NIH 1ZIAES103340 (RES)). The GISAID database was used to access SARS-CoV-2 sequence information; see Supplemental File 5 for table acknowledging the contributions of submitting and originating laboratories.

The content is solely the responsibility of the authors and does not necessarily represent the official views of the NIH.

## Author Contributions

IMW, MNF, and RES conceived and wrote the manuscript. IMW, MNF, JLL, TR, and RES edited and revised the manuscript. IMW, MNF, JLL, and TR created figures. JLL and TR performed all bioinformatics analyses. IMW and MNF purified proteins. IMW performed assays.

## Abbreviations

COVID-19: Coronavirus Disease 2019
SARS-CoV-2: Severe Acute Respiratory Syndrome Coronavirus 2
Nsp15: Non-structural protein 15
RNA: ribonucleic acid
MERS-CoV: Middle Eastern Respiratory Syndrome Coronavirus
SARS-CoV-1: Severe Acute Respiratory Syndrome Coronavirus 1
pp1a: polyprotein 1a
pp1ab: polyprotein 1ab
nsps: Non-structural proteins
RTC: Replication Transcription Complex
dsRNA: double stranded RNA
MDA5: melanoma differentiation-associated protein 5
NTD: N-terminal domain
MD: Middle domain
EndoU: endonuclease domain
GISAID: Global initiative on sharing all influenza data
WT: wild-type
IPTG: Isopropyl β-d-1-thiogalactopyranoside
β-ME: 2-mercaptoethanol
EDTA: Ethylenediamine tetraacetic acid
TEV: Tobacco Etch Virus cleavage site
PMSF: phenylmethylsulfonyl fluoride
SEC: size exclusion chromatography
FRET: fluorescence resonance energy transfer
FI: fluorescein
TAMRA: Carboxytetramethylrhodamine
PAGE: polyacrylamide gel electrophoresis

## Supplemental Information

Supplemental Tables 1-2

Supplemental Figures 1-5

Supplemental Files 1-5

## Legend for Supplemental Files

**Supplemental File 1.** Original GISAID download and analysis, June 15, 2021. See ReadMe tab in spreadsheet for additional information.

**Supplemental File 2.** Feb 20, 2022 GISAID download and analysis. See ReadMe tab in spreadsheet for additional information.

**Supplemental File 3**. Identification of the earliest date of origin of each of the non-synonymous Nsp15 substitutions.

GISAID accession IDs for all Nsp15 proteins that contain each substitution of interest were used to query all of the fasta headers for each Nsp15 protein in GISAID downloaded on 02/15/2022. Each worksheet is a compilation of these headers for a specific substitution of interest, sorted by the oldest date. Each column in each worksheet is a different element in each of the fasta headers for these sequences. The column headers are left in the GISAID format dictated by their upload webpage (https://www.epicov.org/epi3/frontend#5045a9). Column YYY-MM-DD is the collection date for each genome and is the only date we can attach to these proteins. Based on these dates we can estimate the date and location of each of these substitutions. As some of the dates are incomplete, we are basing the date of origin only on dates that were completely provided by submitters.

**Supplemental File 4**. Raw data for the data in Supplemental Table 2.

**Supplemental File 5**. Table acknowledging the GISAID contributions of submitting and originating laboratories.

**Supplemental Table 1:** Plasmid Constructs used in this study. All constructs generated by Genscript (Piscataway, NJ).

| **Construct** | **Mutation Location**<br>**(domain level)** | **First created** |
| --- | --- | --- |
| WT-Nsp15 (6xHis-thrombin-TEV/pet14-b) | N/A | P[28] |
| Nsp15 K13N | NTD | This study |
| Nsp15 K13R | NTD | This study |
| Nsp15 G18E | NTD | This study |
| Nsp15 G18R | NTD | This study |
| Nsp15 T34I | NTD | This study |
| Nsp15 A93S | MD | This study |
| Nsp15 T115A | MD | This study |
| Nsp15 V128F | MD | This study |
| Nsp15 D133Y | MD | This study |
| Nsp15 L163F | MD | This study |
| Nsp15 P206S | EndoU | This study |
| Nsp15 R207S | EndoU | This study |
| Nsp15 D220Y | EndoU | This study |
| Nsp15 H235Y | EndoU | This study |
| Nsp15 K260R | EndoU | This study |
| Nsp15 K290N | EndoU | This study |
| Nsp15 W333C | EndoU | This study |

**Supplemental Table 2:**
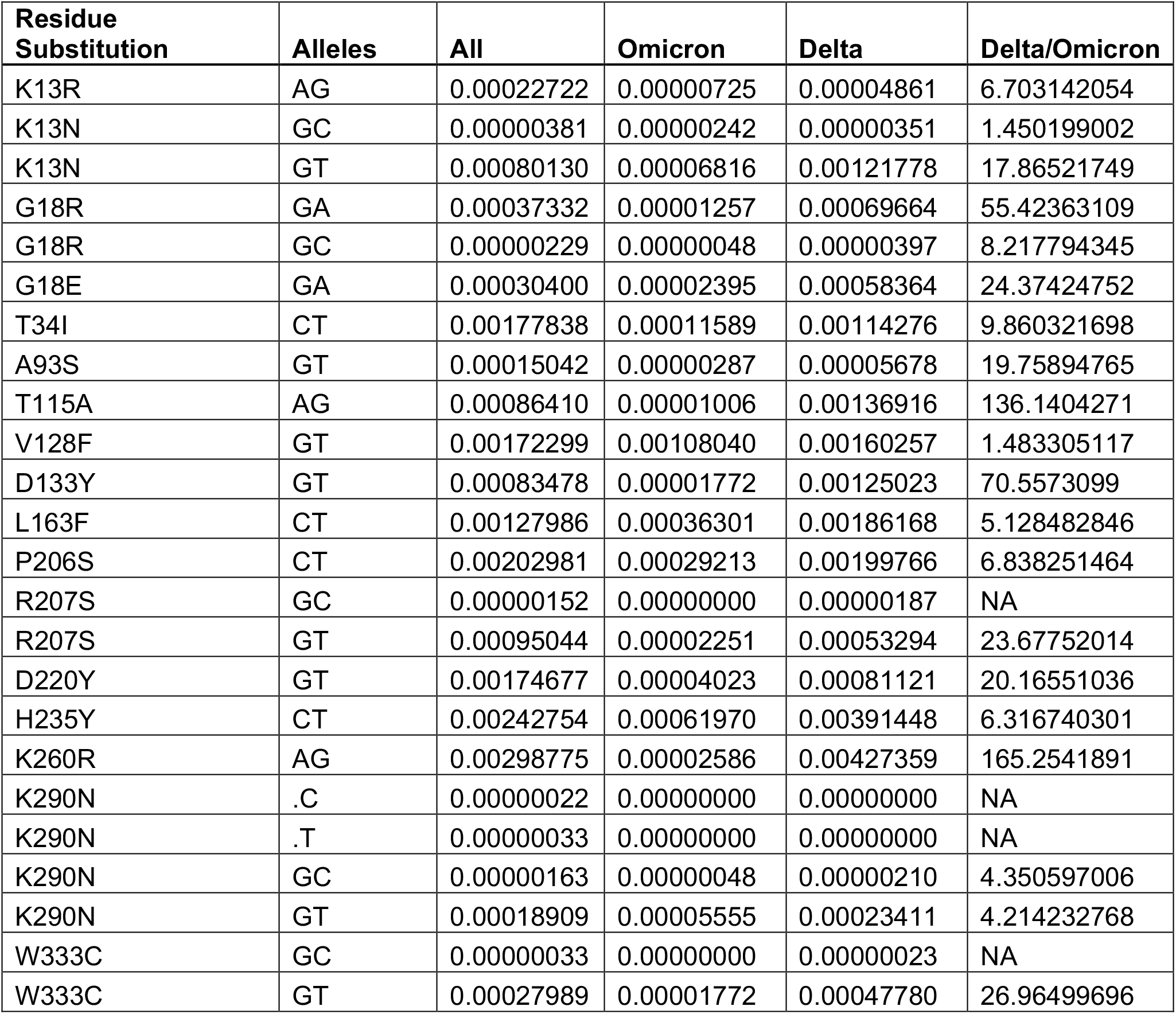
The frequency of occurrence of each non-synonymous Nsp15 substitution in the Delta and Omicron variants of concern. GISAID accession IDs for each of the genomes containing a substitution listed in the Residue Substitution column were used to search the Custom Download page (https://www.epicov.org/epi3/frontend#4fa597) using the Select function. Prior to the search, the “Variant” selector was used to limit the search to either VOC Omicron, VOC Delta, or all variants. All is the frequency of each substitution in all SARS CoV2 genomes in GISAID; Delta is the frequency of each substitution in the Delta subset of SARS CoV2 genomes; Omicron is the frequency of each substitution in the Omicron subset of SARS CoV2 genomes. Delta/Omicron is the ratio of the count of each substitution in each of the two VOC. For raw data see Supplemental File Four.

| Residue Substitution | Alleles | All | Omicron | Delta | Delta/Omicron |
| --- | --- | --- | --- | --- | --- |
| K13R | AG | 0.00022722 | 0.00000725 | 0.00004861 | 6.703142054 |
| K13N | GC | 0.00000381 | 0.00000242 | 0.00000351 | 1.450199002 |
| K13N | GT | 0.00080130 | 0.00006816 | 0.00121778 | 17.86521749 |
| G18R | GA | 0.00037332 | 0.00001257 | 0.00069664 | 55.42363109 |
| G18R | GC | 0.00000229 | 0.00000048 | 0.00000397 | 8.217794345 |
| G18E | GA | 0.00030400 | 0.00002395 | 0.00058364 | 24.37424752 |
| T34I | CT | 0.00177838 | 0.00011589 | 0.00114276 | 9.860321698 |
| A93S | GT | 0.00015042 | 0.00000287 | 0.00005678 | 19.75894765 |
| T115A | AG | 0.00086410 | 0.00001006 | 0.00136916 | 136.1404271 |
| V128F | GT | 0.00172299 | 0.00108040 | 0.00160257 | 1.483305117 |
| D133Y | GT | 0.00083478 | 0.00001772 | 0.00125023 | 70.5573099 |
| L163F | CT | 0.00127986 | 0.00036301 | 0.00186168 | 5.128482846 |
| P206S | CT | 0.00202981 | 0.00029213 | 0.00199766 | 6.838251464 |
| R207S | GC | 0.00000152 | 0.00000000 | 0.00000187 | NA |
| R207S | GT | 0.00095044 | 0.00002251 | 0.00053294 | 23.67752014 |
| D220Y | GT | 0.00174677 | 0.00004023 | 0.00081121 | 20.16551036 |
| H235Y | CT | 0.00242754 | 0.00061970 | 0.00391448 | 6.316740301 |
| K260R | AG | 0.00298775 | 0.00002586 | 0.00427359 | 165.2541891 |
| K290N | .C | 0.00000022 | 0.00000000 | 0.00000000 | NA |
| K290N | .T | 0.00000033 | 0.00000000 | 0.00000000 | NA |
| K290N | GC | 0.00000163 | 0.00000048 | 0.00000210 | 4.350597006 |
| K290N | GT | 0.00018909 | 0.00005555 | 0.00023411 | 4.214232768 |
| W333C | GC | 0.00000033 | 0.00000000 | 0.00000023 | NA |
| W333C | GT | 0.00027989 | 0.00001772 | 0.00047780 | 26.96499696 |

**Supplemental Figure 1:**
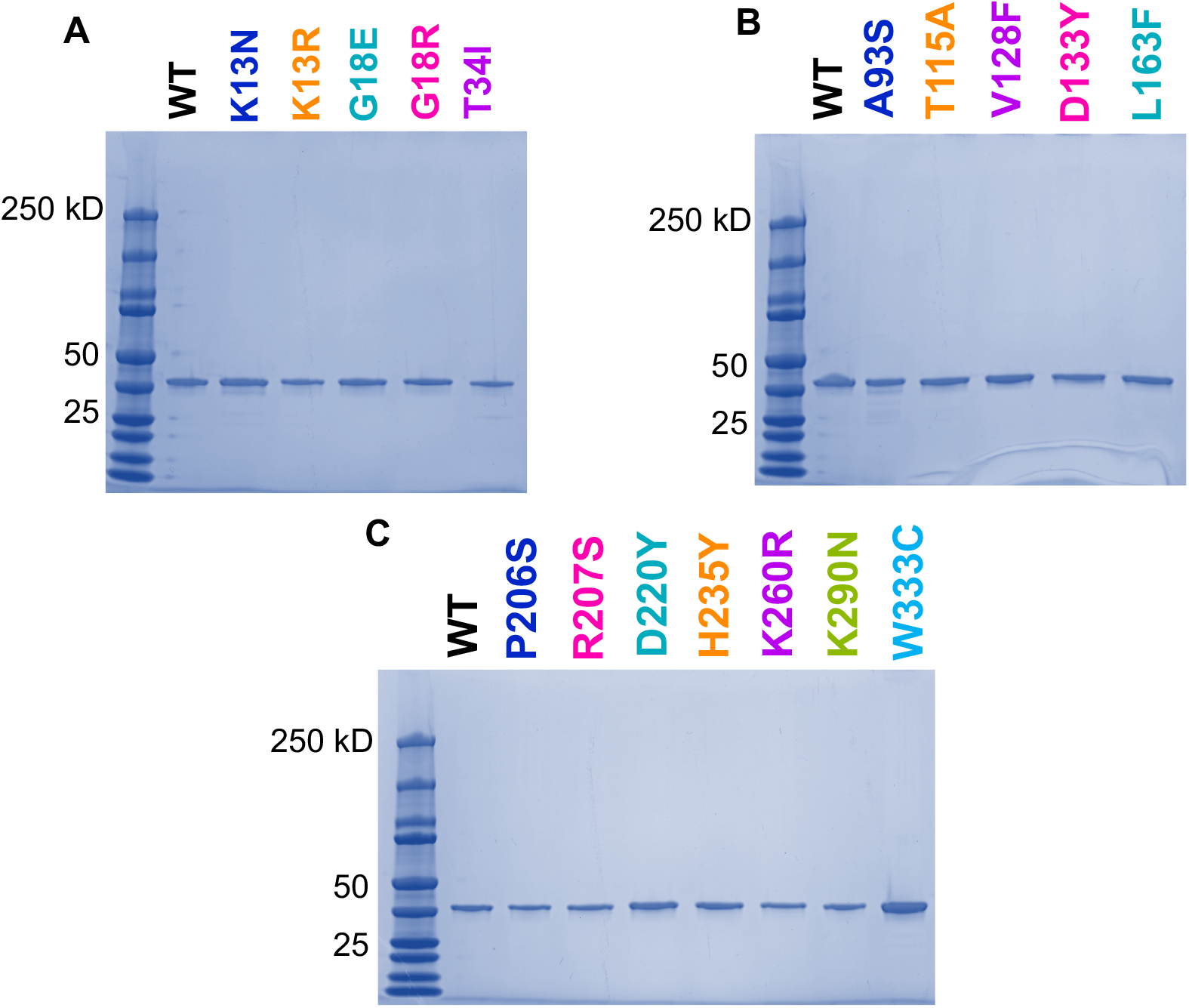
Summary gels following purification of all Nsp15 variants. Fractions corresponding to active hexamer were analyzed by SDS-PAGE and reveal pure protein for NTD **(A)**, MD **(B)**, and EndoU **(C)** variants.

**Supplemental Figure 2:**
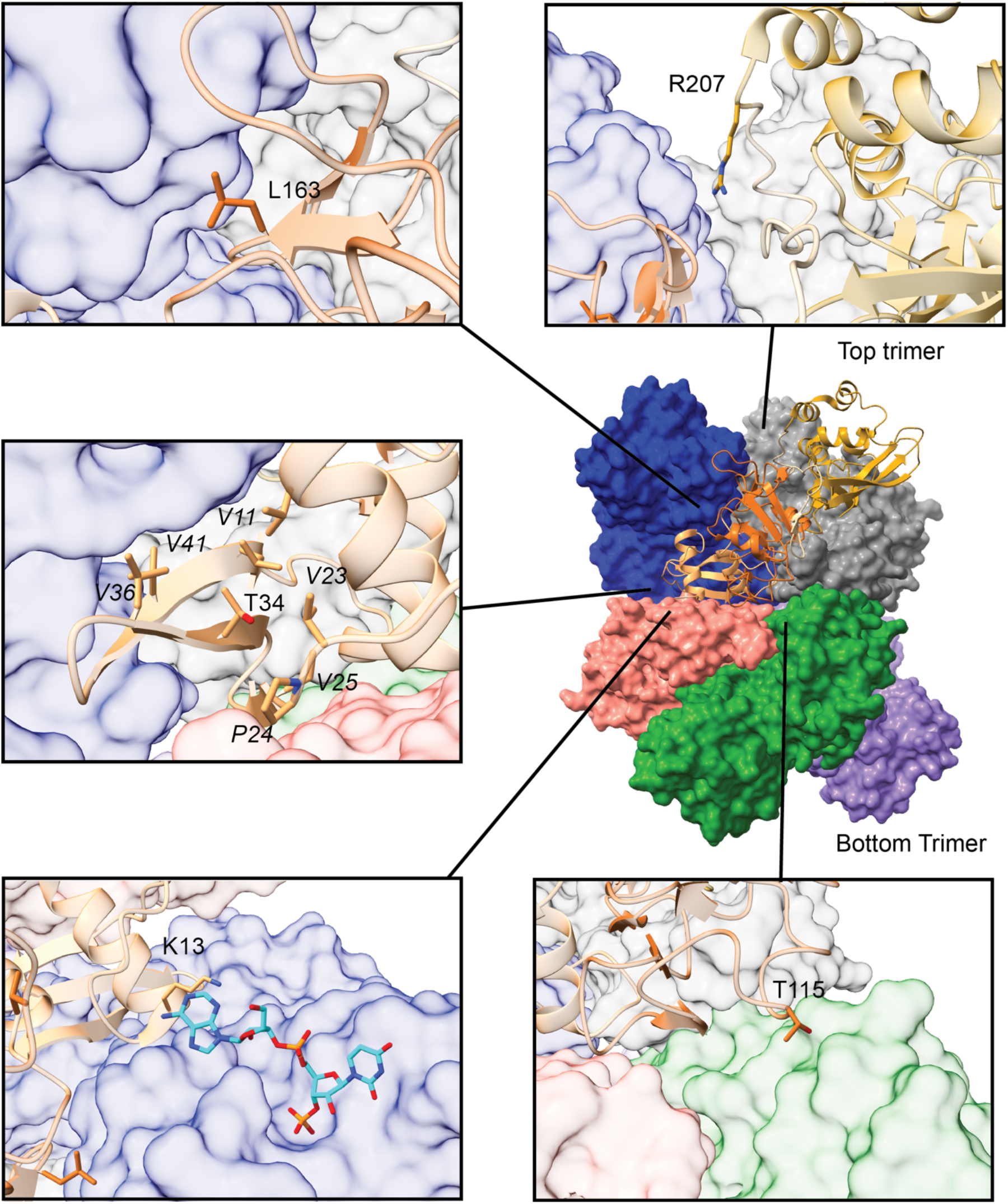
Selected residues involved in hexamer interface interactions. Center right: Nsp15 hexamer (PDB: 7N06). Five protomers shown in surface view (blue, grey, salmon, green, purple); one protomer shown in ribbon view and colored by domain (NTD, tan; MD, orange; EndoU, gold). Boxes depict zoomed in regions with residues of interest. From top right, counter-clockwise: EndoU residue R207, MD residue L163, NTD residue T34 with surrounding core residues labeled in italics, NTD residue K13, and MD residue T115. K13 is shown with a post-cleavage RNA; the 5’ base extends towards the neighboring NTD including K13.

**Supplemental Figure 3:**
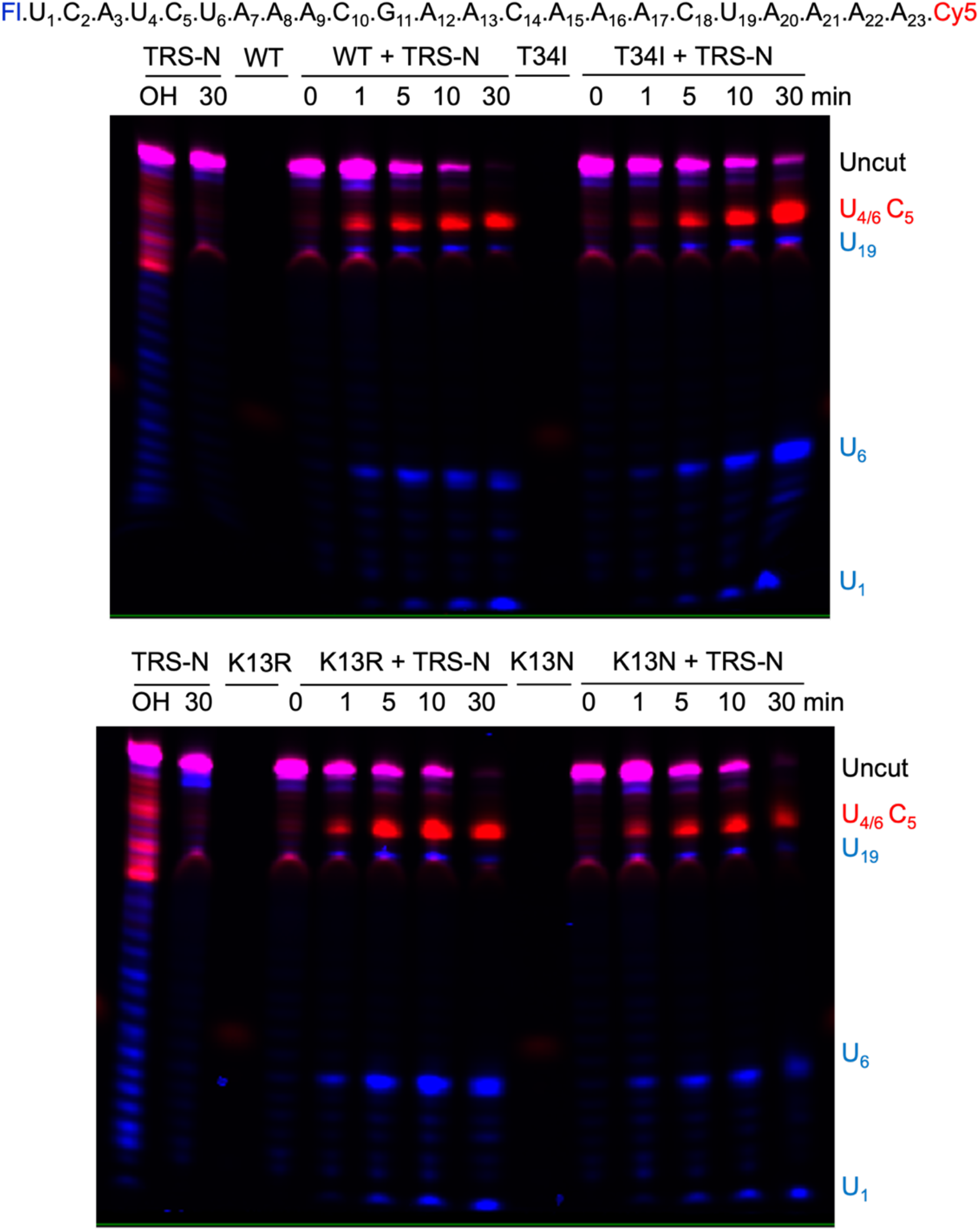
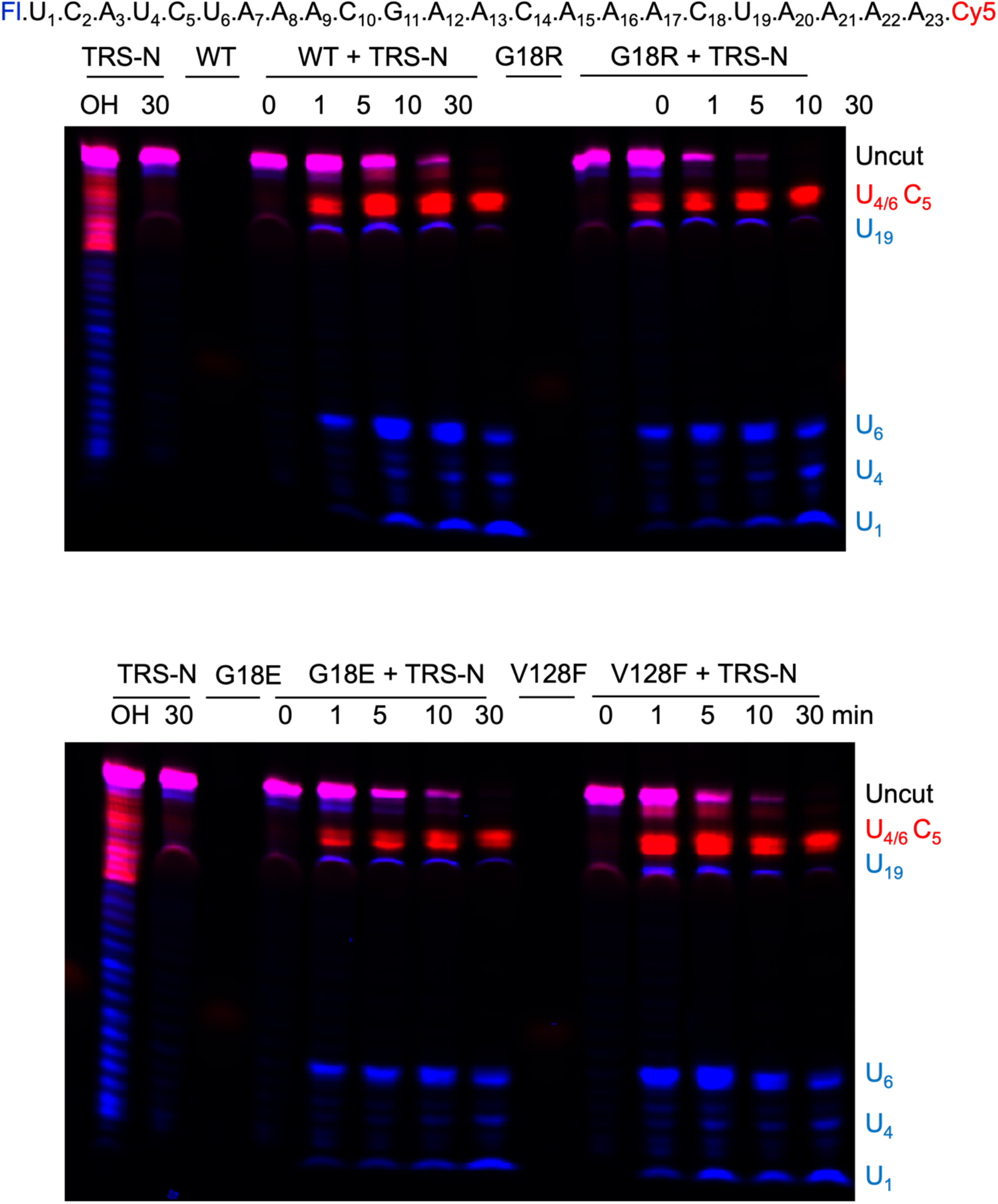
Gel-based endonuclease assays for NTD/MD mutants. The transcriptional regulatory sequence for the nucleocapsid protein (TRS-N) is fluorescently labeled on both end (see labeled sequence at top of each set of gels). A 30-min time course nuclease assay was carried out with WT or Nsp15 NTD/MD variants.

**Supplemental Figure 4:**
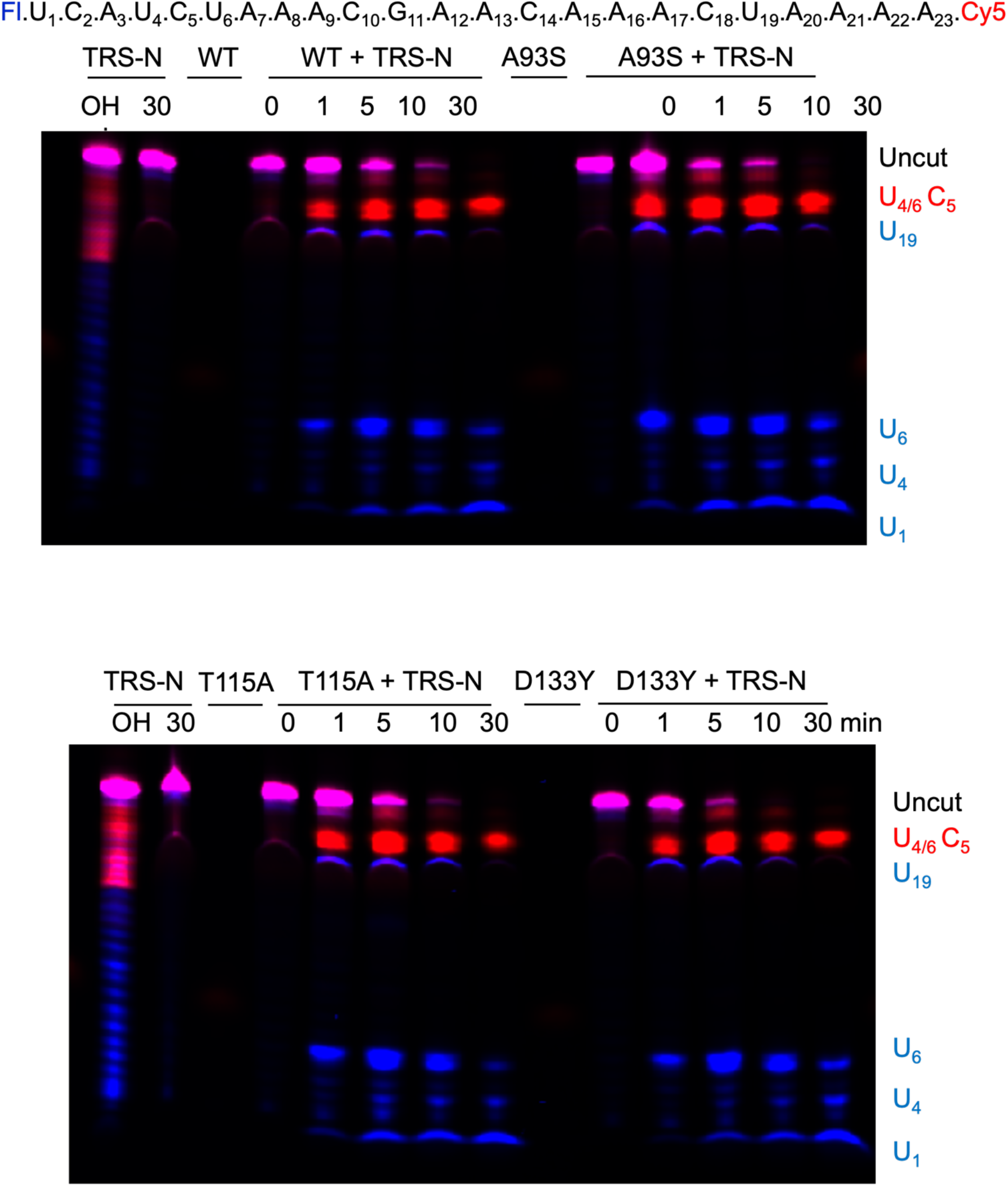
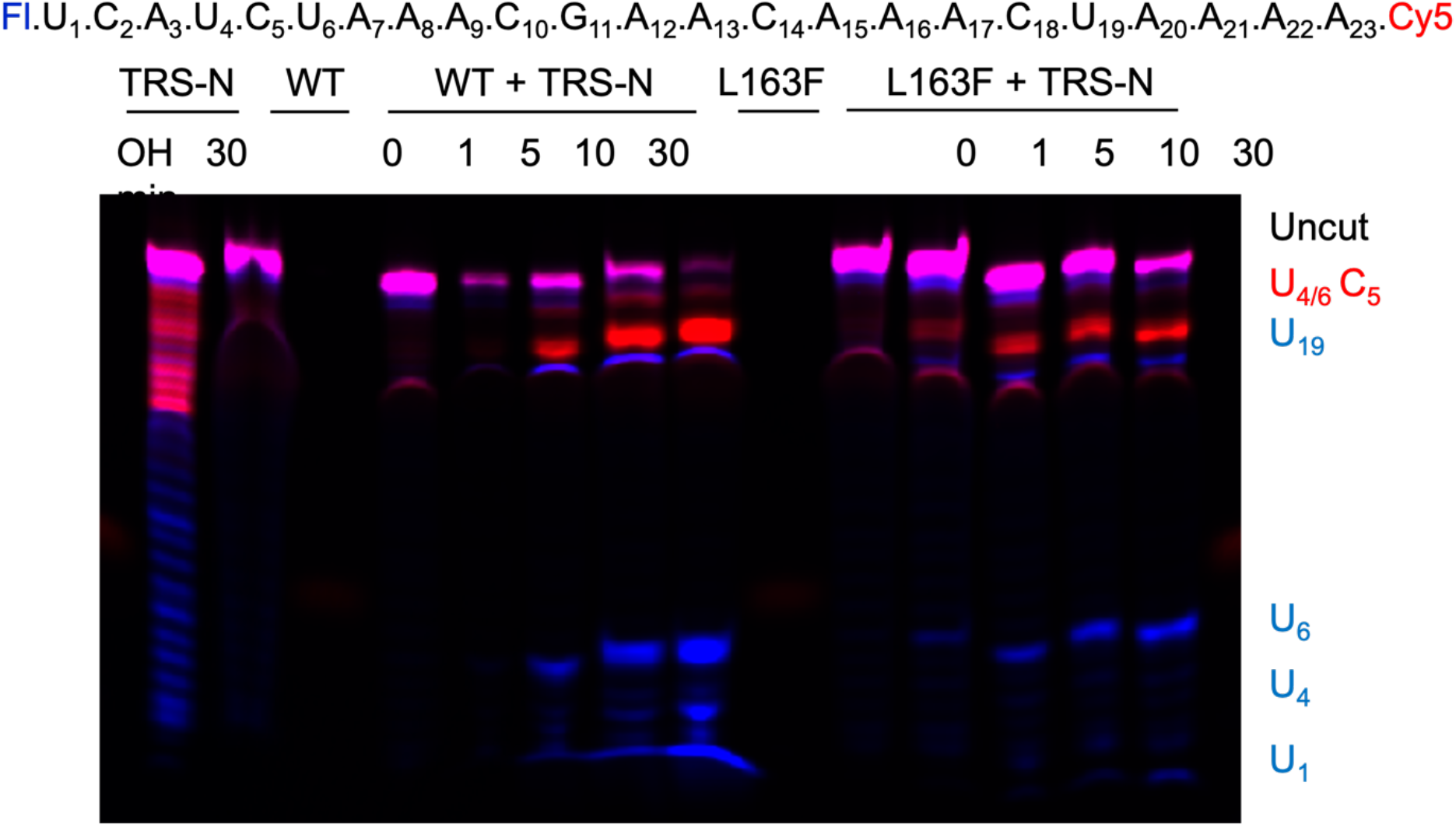
Gel-based endonuclease assays for MD mutants. The transcriptional regulatory sequence for the nucleocapsid protein (TRS-N) is fluorescently labeled on both end (see labeled sequence at top of each set of gels). A 30-min time course nuclease assay was carried out with WT or Nsp15 MD variants.

**Supplemental Figure 5:**
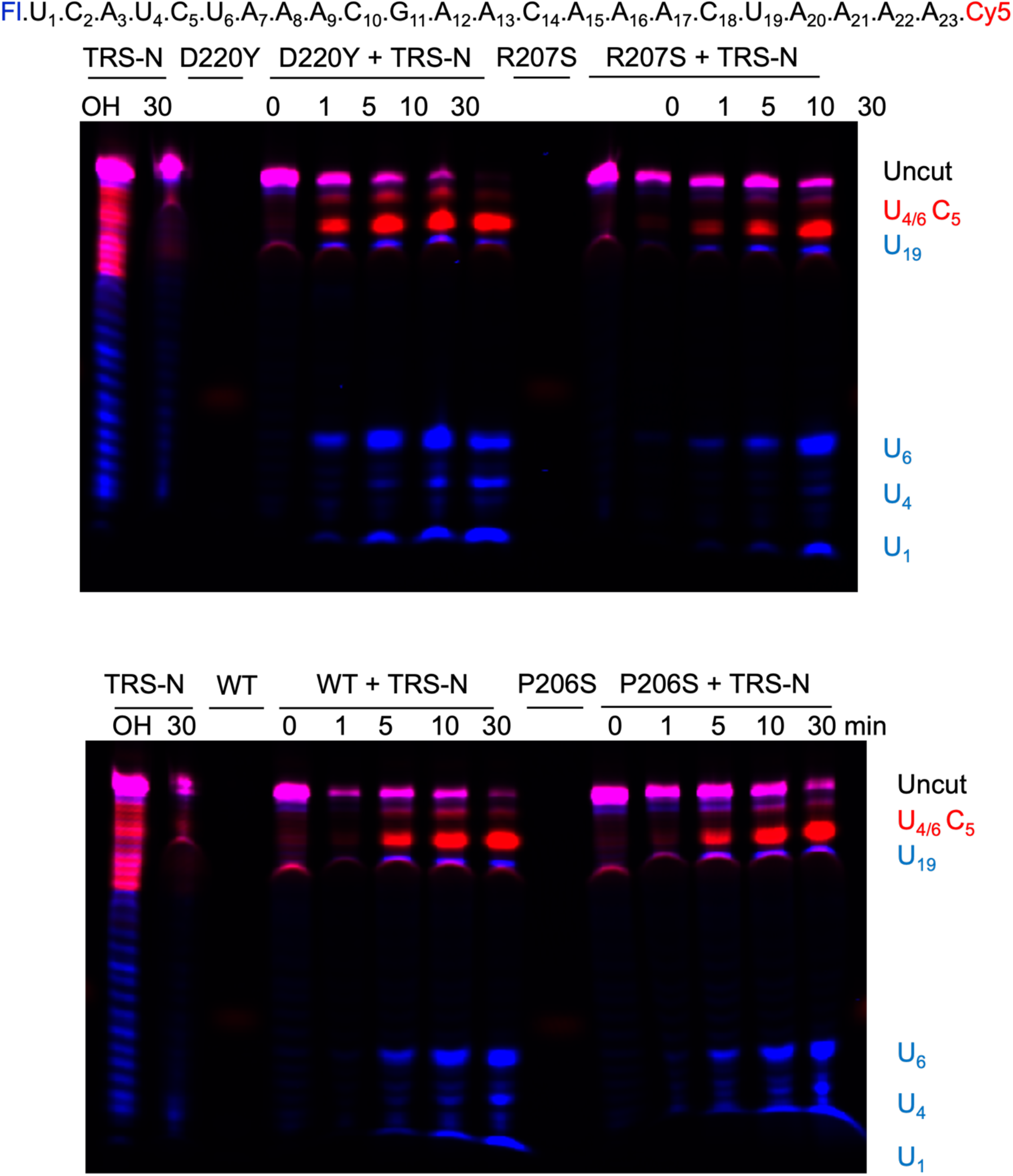
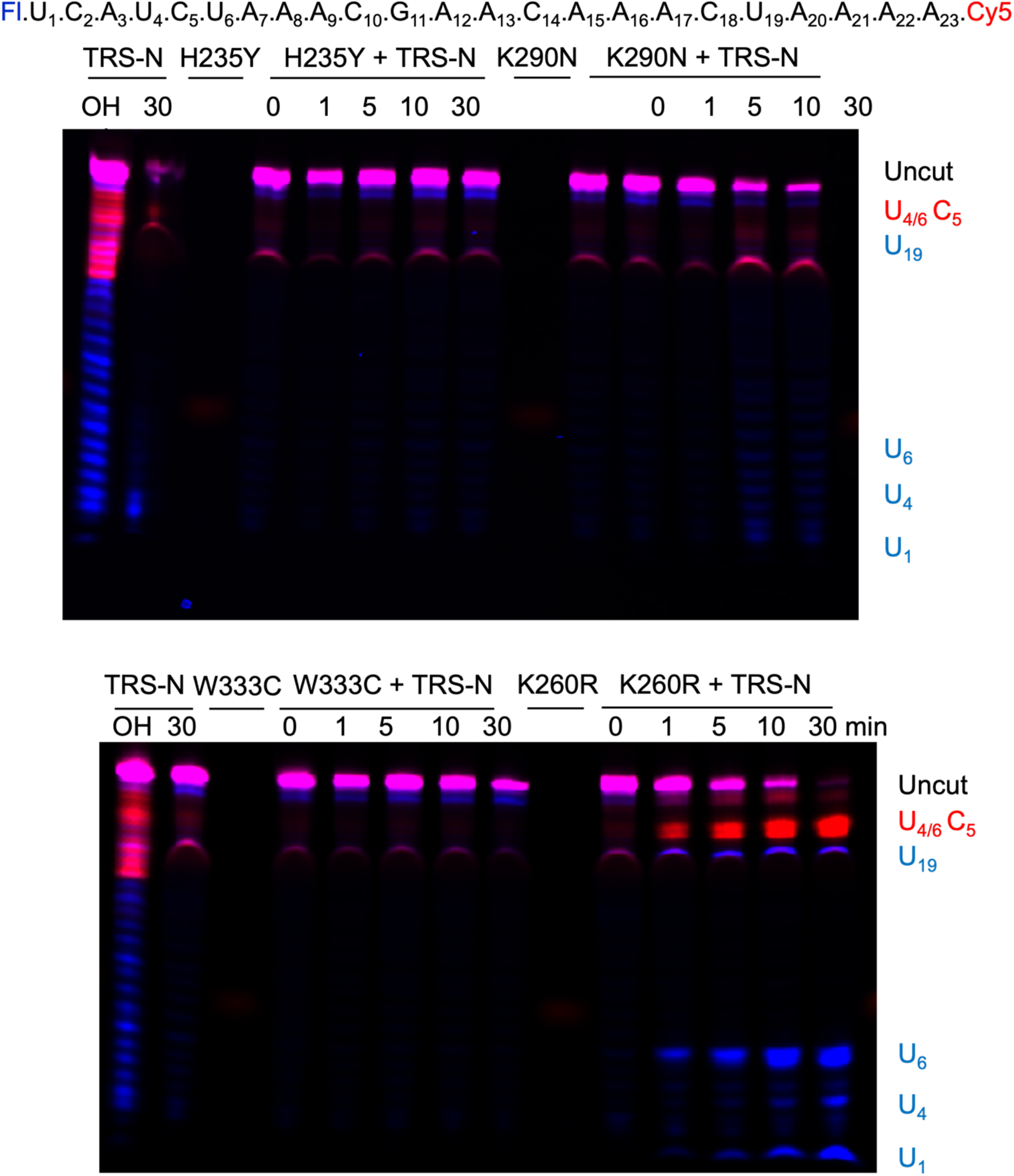
Gel-based endonuclease assays for EndoU mutants. The transcriptional regulatory sequence for the nucleocapsid protein (TRS-N) is fluorescently labeled on both end (see labeled sequence at top of each set of gels). A 30-min time course nuclease assay was carried out with WT or Nsp15 EndoU variants.

